# Distribution modeling of *Lantana camara* and identification of hotspot for future management of the species in the lower Shivalik region of Uttarakhand, India

**DOI:** 10.1101/2020.12.31.424957

**Authors:** Tamali Mondal, Dinesh Bhatt, K. Ramesh

## Abstract

Invasive alien species are one of the most significant threats to biodiversity of the earth ecosystem. Spatial modeling has played a vital role in predicting and mapping the suitable areas of the species habitat for management implementation. *Lantana camara* is one of the ten worst invasive weed enlisted by IUCN., and has its distribution in more than 154000 sq. km area in India. We predicted the distribution of *Lantana camara* using Ecological Niche Modeling (ENM) in MaxEnt software for Rajaji- Corbett landscape in the lower Shivalik region of the state of Uttarakhand, India. The predicted area for the invasion was classified into four categories high, good, medium, and low potential areas. The distribution hotspots were identified using Getis Ord G* method in ArcGIS 10.5 by calculating plot-wise species occurrence. Major hotspot areas under threat of high infestation are Dholkhand, Gohri, Ramgarh, Motichur, and Chilla from Rajaji Tiger Reserve, Palain, Mandal and Maidavan from Kalagarh Tiger Reserve and Bijrani, Dheela, Dhikala, Jhirna and Kalagarh from Corbett Tiger Reserve. This study provides a current status of the invasive species in the landscape with identified areas that need immediate actions to control the spread of the species. The study results can help the policymakers and stakeholders build an efficient and strategically weed management plan.

## Introduction

Migration and colonization are two crucial cycles of life for the survival of any species on Earth (White et al. 2006). Several migrating species were unable to ascertain sustainable populations in new habitats and quickly died out. In alternative cases, they were either incorporated into the existing structure of the ecosystem or were liable for modifying the existing one by competing native competitors and decimating them (Wilcove & Chen 1998).“Invasive species” implies the invaders that have varied effects on native species. About 42 percent of the earth species are at risk due to the expansion of different invasive species. There were overall 120,000 types of plants, animals, and microorganisms that were non-native erstwhile, had invaded the United States, United Kingdom, Australia, South Africa, Brazil, and India (Pimentel et al. 2000). It is presumed that more than 480,000 species had invaded the biological system over the planet earth. In India, a population of 1599 alien plant species belonging to 842 genera in 161 families can be found, which comprises 8.5% of the country’s total vascular flora. (Khuroo et al. 2012; Mandal and Joshi 2014). To control the spread of the invasive species, a zonal control for various invasion levels needs to be in place in the targeted areas (Grice et al. 2011, Terblanche et al. 2016, and Shackleton et al. 2017). To develop that kind of strategy it requires a comprehensive knowledge of a species ecology of invasion, including factors influencing its nearby abundance, spread, and effect. A fundamental survey is needed before structuring such productive methodology which will address (1) the present and the future geographic distribution and abundance of the target species, and (2) it’s status within the invaded range in correlation with the global or local environment under current and future climate (Eckert and Kilawe 2020).

The study here is significantly centred around *Lantana camara*, which used to be an ornamental plant; presently enrolled inside the world’s top ten invasive plant species (Ghisalberti 2000, Richardson and RejManek 2011), and a serious threat to native plants in India (Mungi et al. 2019). The species was introduced as a garden plant by the Britishers at National Botanical Garden, Kolkata, in the year 1807 (Nanjappa et al. 2005; Bhagwat et al. 2012). Lantana is a medium-sized woody bush, local to central and South America, used to have multiple stems (Sundaram and Hiremath 2012). The synthetic breed, which has been used for plant improvement in Europe as early as the sixteenth century, now exists in various structures or assortments worldwide. It has been cultivated for more than 300 years and now has many cultivars and hybrids. These hybrids generally fall under the L. camara complex. Cultivars can be recognized morphologically (because of variety in blossom size, shape, and shading; leaf size, bushiness, and shading; stem prickliness), physiologically (by analyzing the variation in development rates, poisonousness to domesticated animals), and by their chromosome number and DNA content (Binggeli, 1999).

*Lantana’s* diverse and immense geographic distribution mirrors its wide natural resistances. However the distribution of *Lantana* is limited by its inability to sustain under thick, unblemished shades of taller local timberland species (IUCN). *Lantana* cannot sustain itself in frigid temperatures (Corlett 1992, Thinley 2020), to saline soils. It is inclined to spoil in boggy or hydromorphic soils; it has never been acquainted with certain islands; low precipitation, and coralline soils with helpless water-holding limits; and high rate of tropical typhoons (Thaman 1974, Day et al. 2003). Whereas, It occurs on an assortment of soil types and in various natural surroundings. In open unshaded circumstances, for example, wastelands, rainforest edges, seashore fronts and regenerating woods from fire or logging it, for the most part, develop best. Likewise, disturbed regions, for example, streets, railroad tracks, and channels, were suitable for the species (Thaman 1974; Winder and Harley 1983; Thakur et al. 1992; Munir 1996; Day et al. 2003). Lantana was found on the edges of the rainforest instead of invading new territories (Humphries and Stanton 1992; Day et al. 2003).

A series of widely accepted modeling approaches are developed to check invasive species’ ecological niches of invasive species to know their current stages of invasion and potential distribution in space or time (Guisan and Thuiller 2005; Elith and Leathwick 2009). Two modeling approaches which stand out amongst them are mechanistic models and statistical correlative model. Mechanistic models require collection and validation of massive amounts of physiological data because it deals with functional traits of the species. The statistical correlative model, geographic species occurrence and preexisting environmental data are the keys (Kearney and Porter 2010). There are two prevailing techniques for statistical modeling approach: ordination, which relies on direct observations, and species distribution modeling (also referred to as ecological niche modeling), which depends on predictions based on knowledge of the preferred environment of the species (Peterson and Vieglais 2001; Hirzel et al. 2002; Phillips et al. 2004). Both are viewed as integral to the next. They rely on the specialty idea and depend on the niche concept and relate biotic and abiotic conditions to the likelihood of a species being available in a particular area.

For prioritization and management of invasive species, it is vital to identify the regions at high infestation conditions. The term ‘invasion hotspot’ (O’Donnell et al. 2012) was used earlier for areas with a high potential for colonization by diverse alien invasive species. Due to anthropogenic pressure like biodiversity loss, agricultural expansion, shifting cultivation, and urbanization, the forest cover declines gradually. These disturbances help to create space for invasion (Shea and Chesson 2002; Gravuer et al. 2008; Catford et al. 2011; Bellard et al. 2014). The concept of availability of “vacant niches” explains that the ecological requirements (soil nutrients, specific climatic conditions, required habitat) of the invasive species in these empty spaces get fulfilled naturally (Bellard et al. 2014; Moles et al. 2008; Lekevičius 2009; Denslow JS 2003).

The main objective of this work was to identify potential areas for *Lantana* occurrences and define ‘invasive hotspots’ for management prioritization of the species in the Rajaji Corbett Landscape established in the lower Shiwalik Himalayan plateau on the Gangetic Plain. We have identified lantana hotspots in the study area by combining various biotic and abiotic parameters and data collected through direct observation. To identify potential Lantana habitats, an ecological niche model (ENM) was performed using MaxEnt 3.4. This study adopted a combination of species distribution modeling and hotspot analysis to identify the high-density lantana infestation site, which is crucial for a functional management plan within the area.

### Study Area

This study was carried out in parts of the Terai Arc Landscape of the lower Shiwalik Himalayas, under the jurisdiction of Haridwar and Pauri Garhwal district of Uttarakhand (Dhaundiyal 1997). The landscape covers an area of about 2,363.48 sq. km of Bhabar tracts bounded by the rivers Sharda and Yamuna and has about 36% forest cover (Johnsingh et al. 2004). The elevation range of the study area varies from 185m to 1352m asl. The climate is characteristic of the Shivalik landscape, with temperatures ranging up to 42°C in summer and 3-5°C in winter. Rainfall ranges from 1,253 to 2,053mm. The forest types of this region categorized into North Indian Deciduous Forest (which includes Moist Siwalik Sal forest), West Gangetic moist mixed deciduous forest, Northern Tropical dry mixed deciduous forest, Dry grassland, Scrub, and Dry bamboo bakes (Champion and Seth 1968). Apart from the naturally occurring forests, plantations of several species viz., *Eucalyptus spp*, *Tectona grandis, Halophragma adenophyllum, Acaciacatechu, Dalbergia sissoo*, and *Ailanthus excels*are common in this area (Johnsingh et al. 2005). The landscape has a diverse range of grassland and wetlands dominated by Graminoid species of *Saccharum sp*, *Narenga sp, Imperata cylindrical*, and *Typha sp* (Mathur 2000). Sal (*Shorea robusta*) is the most dominant species in the area, and it has floral elements of peninsular India. The region also has floral species closely related to those found in the eastern Himalayas or the Western Ghats like *Schefflera venulosa, Diospyros embryopteris, Phoebe lanceolota, Wallichia densiflora* and *Bischoffia javanica*. Two endemic species found in this region are *Eremostachys superba* and *Catamixis baccharoides*.

The landscape includes several forest divisions and protected areas under the administration of 20 Forest Divisions, among which Rajaji (Rajaji TR) and Jim Corbett Tiger Reserves (Corbett TR) are the major ones (Figure 1). This landscape is primarily dominated by dry deciduous forest, but the northeast part of it (belongs to Kalagarh Tiger Reserve) holds the high elevated (1200-1300m) areas where vegetation community changes. Shrubs like Curry patta (*Murraya koenigii*), Rohini (*Mallotus philippensis*), Kadu (scientific name not available), Kakra (scientific name not available) were the dominant species found here along the roads apart from *Lantana*. High tree density is relatively abundant within the core areas on this part of the landscape. The main connectivity of this area depends on a two-way lane that connects Kotdwar to Ranikhet. This road also acts as the only barrier between the reserve boundary and the villages situated around the Kalagarh Tiger Reserve. This road connectivity in this part is also made by demolishing the hilly terrains, which made these entire roadside hilly edges bare and potential habitat for invasion. During our survey, we observed very high infestation of Lantana in these areas.

**Figure 1.**
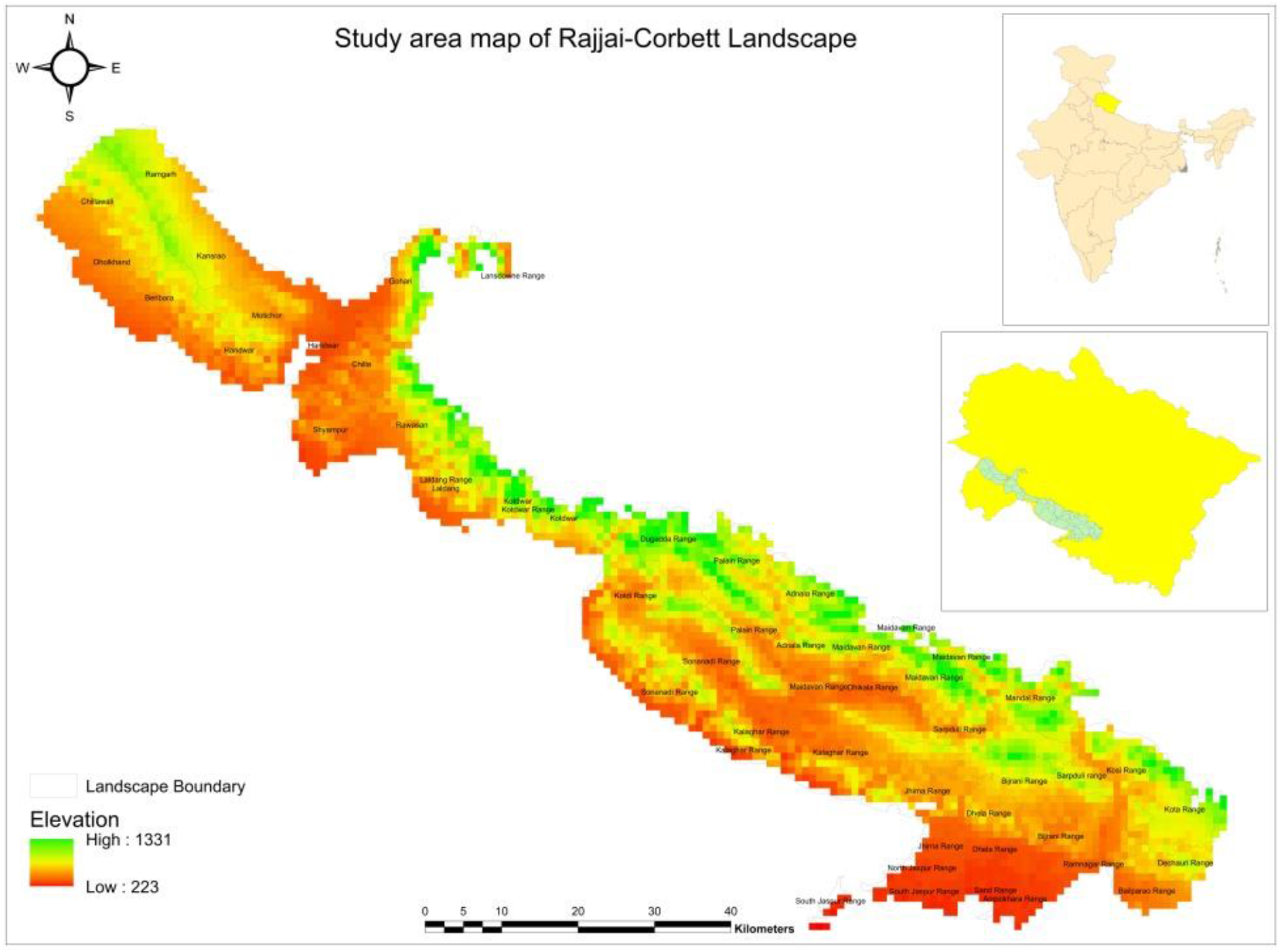
Map of the Study area

**Figure 2:**
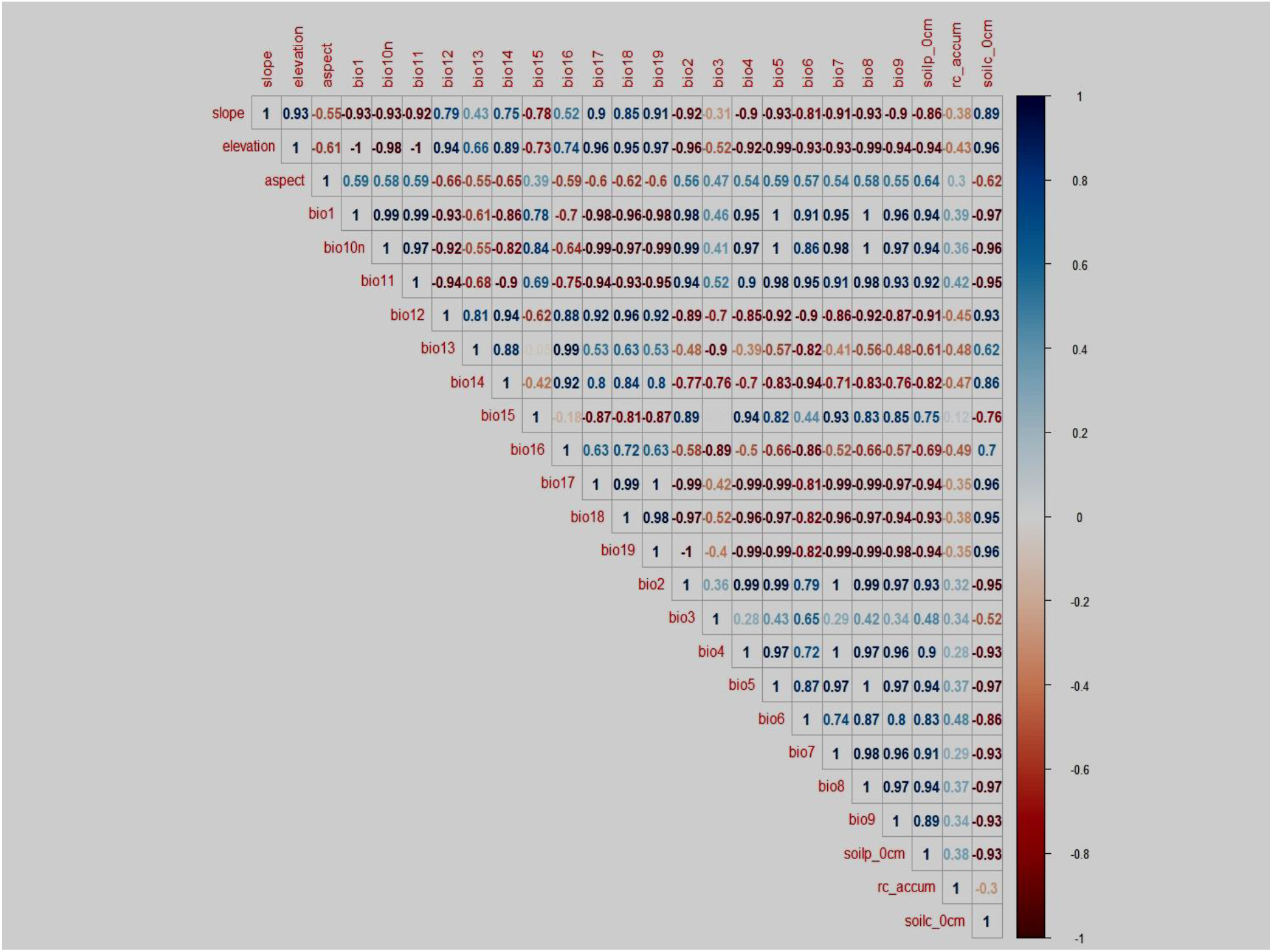
Correlation coefficient between the variables used

**Figure 3:**
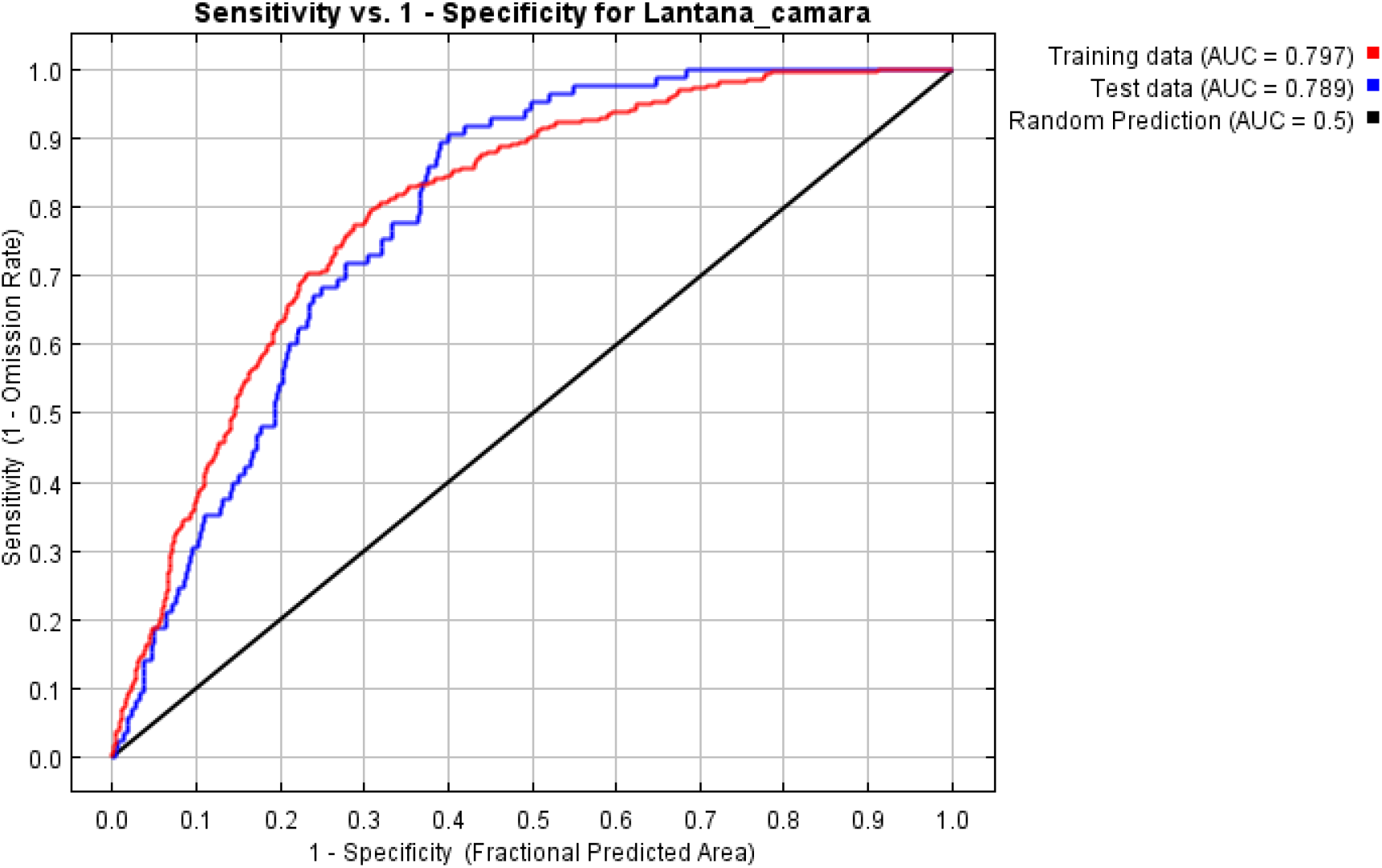
ROC curve of sensitivity versus specificity for *Lantana camara*

## Methodology

### Data collection

Studies have indicated that the threshold for invasive occurrence between successive plots is arbitrary. Research in East Africa used an interval of 25 km for the survey (Wabuyele et al. 2014), while another research conducted in Arunachal Pradesh, India took an interval of 5–30 km (Kosaka et al. 2010). Research by ICIMOD Nepal (2018) in Kailash Sacred Landscape used 5–10 km of interval for the survey. As the region falls in the Shivalik Himalaya, below the middle Himalayas, the elevation of the study area decreases gradually; the climate also influences the shorter distances in the shift in vegetation. It is not just the height but also dimension and other topographical variables that change dramatically within a short distance in the foothills and mountains. Shorter intervals are therefore necessary to properly reflect microhabitats (Thapa et al. 2018). Acknowledging the geophysical variations induced by landscape features, we opted for an interval of 1 km for this study. The research area consists of several protected areas for which a proper permit was obtained.

Road networks frequently act as the carrier for dispersal of invasive plant species because of significant distance dispersal of propagules by vehicles (von der Lippe and Kowarik 2007) just as the street, borderlines being appropriate for colonization by non-native plant species (Christen and Matlack 2006; Johnston and Johnston 2004). Therefore, a roadside survey at the beat level was conducted for rapid assessment of the distribution of *Lantana camara* at the landscape level (Wabuyele et al. 2014; Shrestha 2014; Kosaka et al. 2020). Since landscape-level sampling itself is a costly and time-consuming procedure, instead of a single highway, we used several networks of roads and trails at the beat level within the protected areas for mapping. Other than road dispersal the spread of *Lantana* also depends on the dispersal of its seeds by frugivores (Ramaswami et al. 2016). So, presence location inside the dense forest patches is also considered.

From the starting point of the survey, field plots of 10×10m were laid on the alternate sides of the road to record the presence/absence of *Lantana camara*. The current distribution of *Lantana* in the landscape was sampled using a geographical positioning system (GPS) during November 2019-January, 2020. We recorded the species’-ordinates of the species using a handheld GPS (Garmin trex 30x) along roads, river beds, settlements, forest gaps, and fire-prone areas. We selected parameters for studying the plots by reviewing published literature, thus selecting tree cover, shrub cover, tree and shrub density, distance from nearest fire locations, presence of any anthropogenic activity (road, human, trail, logging, lopping, grazing), and percent area covered by a *Lantana* patch (Thapa et al. 2018; Thinley 2020). For the occurrence of *Lantana* (presence/absence), a total of 744 plots were investigated. Additionally, some opportunistic observations were made to capture records and locations of the maximum possible distribution of invasive plants other than *Lantana*. Information on topographic characteristics like elevation, slope, aspect, curvature (Kumar et al. 2009), size, and pattern of different land use (Vila et al. 2011), were recorded as these are some key variables in the distribution and invasion of *Lantana*.

### Environmental and Bioclimatic Data

The distribution of plants in many ways regulates by different climatic conditions (Grace 1987; Luoto 2007; Vicente et al 2010). Information of bioclimatic variables for the study area (19 bioclim layers) was obtained at ~1km² resolution from Worldclim Climate Database, Version 1 (http://www.worldclim.org/). Data for topography was extracted from CGIAR-CSI (Consortium for Spatial Information) data set. The files were mosaicked to a desirable extent, and topographic index (elevation, slope, aspect, flow direction, and flow accumulation) was computed (using ArcGIS 10.5).

The normalized difference vegetation index (NDVI) 2019 and was extracted from Landsat 8 OLI data set. The data set is freely available on the USGS website (www.usgs.gov).

Data on soil characteristics (soil ph, soil carbon content, soil water content, soil texture) obtained from the Open Land Map (openlandmap.org). Since the species exhibits significant associations with land use and are widespread in an open-canopy forest, near water, and along highways, we also acquired data on land use from Globe cover to aid in subsequent analysis (http://geoserver.isciences.com). All environmental variables were resampled to 1 km resolution. The Multi-collinearity of the environmental variables was assessed using the Pearson correlation coefficient.

### Modeling Approach

*Lantana’s* distribution was modeled using maximum entropy ecological niche modeling (MaxEnt) software 3.3.3k (http://www.cs.princeton.edu/~schapire/maxent/). MaxEnt generates an estimate of the species’ likelihood of presence varying from 0 to 1, i.e., from the lowest to the highest distribution probability. The predictor variables used for the modeling were reduced to 16 after eliminating the high correlated values to avoid biases and considered ecologically meaningful for *Lantana’s* sustenance in its required climate.

These variables include nine climatic variables, i.e., mean diurnal range (mean of monthly (max temp - min temp)), isothermality, min temperature of the coldest month, mean temperature of wettest quarter, precipitation of wettest month, precipitation seasonality (coefficient of variation), precipitation of driest quarter, precipitation of warmest quarter, precipitation of coldest quarter, three topographic variables, i.e. slope, aspect, elevation, three environmental variables, i.e. present land use land cover (lulc) of the study area, soil ph. quantity in 0 cm depth, the soil carbon content in 0 cm depth, and one hydrographic variable, i.e., flow accumulation.

There are three output options available in MaxEnt modeling, i.e., raw, cumulative, and logistic. The default output is logistic. Models were run in all of the outputs as mentioned earlier to notice any variation is there or not. Finally, the logistic output is selected, and the model was run following convergence criteria set at 0.01, maximum iterations to 5000 with default values of feature types such as linear, quadratic, product, categorical, threshold, and hinge.

To assess the model’s statistical significance a subset of 75% data was used to trial run the model and 25% of the data was used to validate the model (Thapa et al. 2018). The multiplier for regularization was set at 1. Output was obtained in a logistic format that gives a likelihood of occurrence ranging from 0-1 percent. The AUC values ranged between 0 and 1. Values between 0.2–0.5 were considered weak, 0.5–0.7 moderate, and > 0.7 as high when validating the model’s findings (Adhikari et al. 2015, Thapa et al. 2018, Priyanka and Joshi 2013). To determine the value of significant variables (Yang et al. 2014, IGES, ICIMOD. 2013), the jackknife technique, alternately known as ‘leave one out’, was followed (Phillips 2006; Baldwin 2009). The output was divided into four classes based on the prediction value of the distribution analysis (i) Least (ii) Moderate (iii) Good and (iv) High potential for invasion distribution, and the map was prepared in ArcMap 10.5.

We also performed hotspot analysis based on the ensemble prediction with Getis-Ord Gi (Getis and Ord 1992) statistic using the corresponding tool in Spatial Statistics Tools of ArcGIS 10.5 (ESRI, Redlands, CA, USA). This statistical analysis decides whether the number of species presence at a defined location (GPS points) against a neighborhood position is higher than expected at random relative to the mean for a given attribute in the entire study area. This technique helped to better reveal areas with clustered higher susceptibility (hotspots) or lower susceptibility (coldspots) for the minimum establishment and spread. The output comes with z and p-value that help in the interpretation of the result. A high z and small p-value for a feature indicate a spatial clustering of high value. The higher the z score, the more intense the clustering. To determine the effects of anthropogenic disturbance on *Lantana*’s distribution buffer analysis was performed using 1km and 5km buffer circle around each hotspot sites to calculate the distance between the hotspot zones to the nearest anthropogenic activities (road, raw, settelement) in ArcMap 10.5.

## Results

The simulation accuracy of the model (the precision of the model) was verified through ROC plot. Specifically, the ROC curve (Fielding and Bell 1977) is used to test the model’s simulation accuracy (Hanley and McNeil 1982). The area under the curve (AUC) value shows the model’s predictive precision. In the present analysis, the AUC of the built model was 0.797 and 0.789, for training and test data respectively, based on the possible climatic and environmental factors influencing *Lantana* distribution (Priyanka and Joshi 2013; Philips 2006).

The relative contributions of the variables in the model showed precipitation of the warmest quarter (bio 18) is the most influential (19.1 %), followed by mean temperature of the wettest quarter (bio8) (18.2%), precipitation of driest quarter (bio17) (15.4%) and river accumulation (10.8). Mean temperature of coldest month, slope, precipitation of wettest month and soil carbon content resulted as the second most influential environmental parameters followed by mean diurnal range, isothermality, precipitation of coldest quarter, precipitation of seasonality, aspect, elevation and soil ph.

The outputs were imported to ArcGIS and converted to a tiff raster format for further analysis. The output maps of the distribution of *Lantana* generated by MaxEnt were classified into four classes viz., <0.20, 0.02-0.40, 0.04-0.60, and >0.60 as least, moderate, good, and high potential areas for Lantana invasion (Priyanka and Joshi 2013; Thapa et al. 2018). *Lantana camara* was found to have invaded Ramgarh, Dholkhand, Beribara, Motichur, Haridwar, Gohri, Chilla, Rawasn, Laldhang, Kotdi, Pakhru, Palain, Kalagarh, Dhikala, Maidavan, Sarpduli, Bijrani, Dhela, and Mandal ranges of the landscape (Fig 4). A significant difference was observed in the habitat niche of *Lantana* within the two tiger reserve. In Rajaji TR, the high elevation areas exhibited low infestation whereas in Corbett the invasion was high in all elevation gradients.

**Figure 4:**
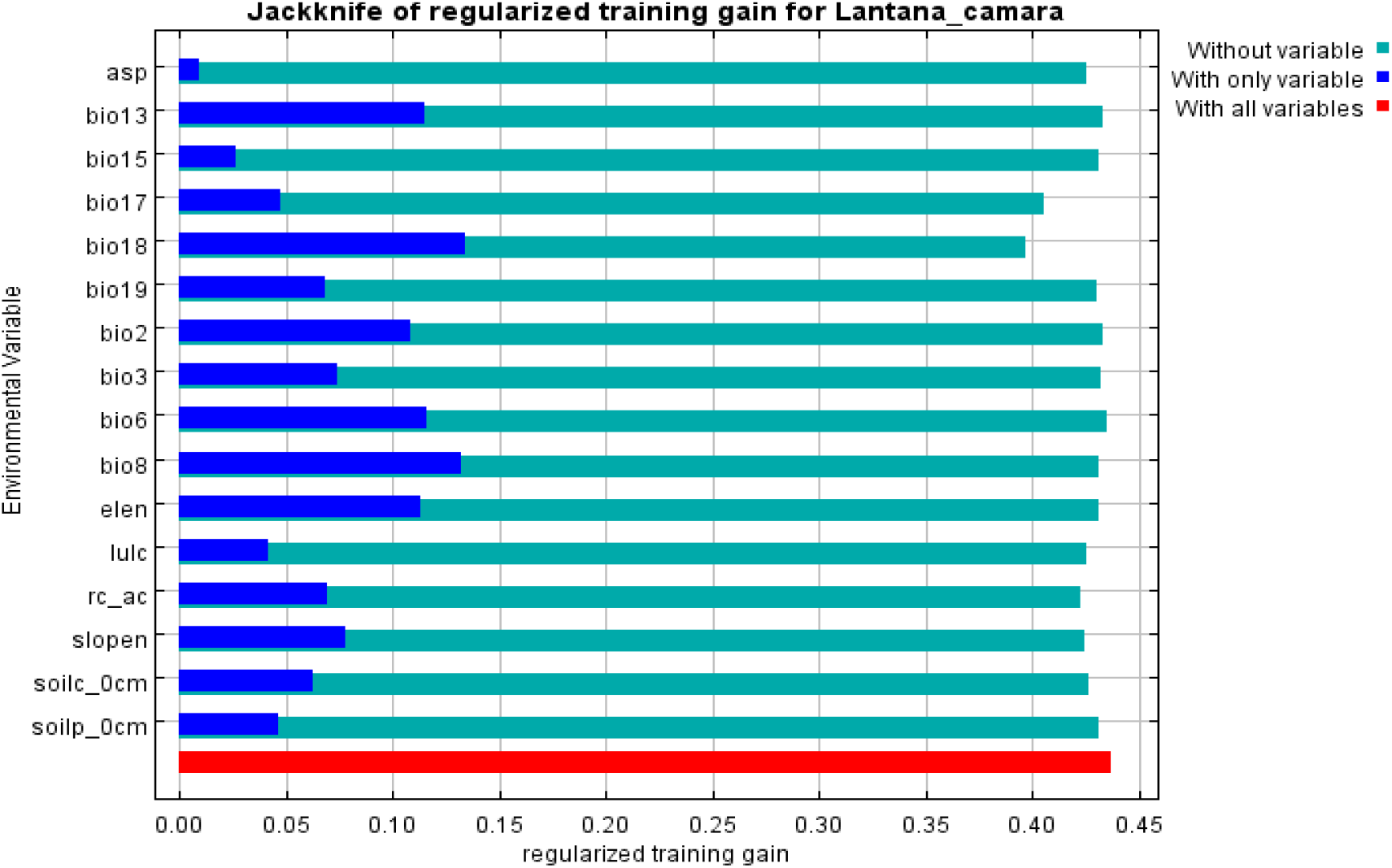
The Jackknife test for evaluating the relative importance of environmental variables for *Lantana camara* in Lower Shiwalik Region Note: ‘asp’ is Aspect, ‘elen’ is elevation, ‘rc_ac’ is flow accumulation, ‘soilc’ is soil carbon content, ‘soilp’ is soil ph.

**Figure 5:**
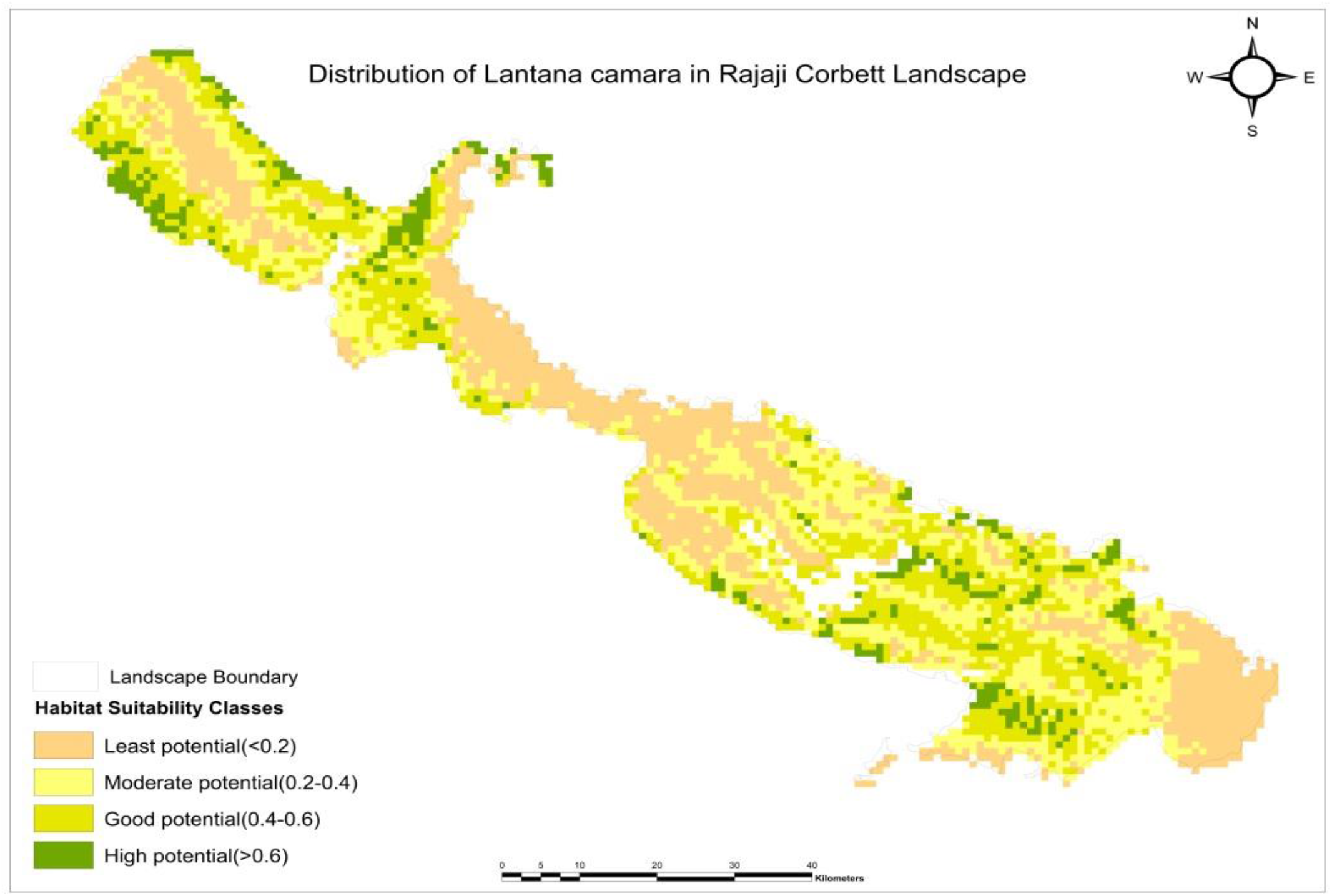
Predicted potential distribution of *Lantana camara* in Rajaji Corbett Landscape

The Hot Spot Analysis based on the statistics Getis-Ord Gi* after selection of the areas showing the highest z-scores (i.e., the two highest confidence intervals, with p = 99% and p = 95%, respectively) highlighted nine ‘hotspots’ of the *Lantana* invasion in the landscape (Ramgarh, Motichur, Dholkahnd, Chilla, Gohri, Palain, Dhikala, Mandal, Maidavan, Jhirna, Dheela, Bijrani, Kalagarh). The hotspot analysis gave a visual interpretation of the specific areas of infestation. As previous studies have recorded, roadside environment, disturbed soil, and dry river beds provide suitable microhabitat for invasive plants; a distance analysis was performed by layering road, river, and 1km buffer areas to each hotspot zone in ArcMap 10.5 (Upadhaya et al. 2019). The output showed each invasion hotspot in the landscape is situated more or less within 1km from either road or river areas.

## Discussion

India holds 712,349 sq. km forested area, among which 154,000 sq. km has been invaded by Lantana, (Mungi et al. 2020). The study by Mungi et al. 2020 also estimated that 40% of the Indian forest is at a high risk of biodiversity loss by Lantana invasion. Numerous studies have already been done and going on to find the best method to control invasive occurrence or spread around the globe. Along with the traditional cut rootstock method, burning, mechanical grubbing, chemical control, and biological control also have failed. Habitat restoration has shown results in very small scale but no records of large scale habitat restoration for *Lantana* eradication are unknown yet (Love et al. 2009).

The species distribution model/ niche model/ habitat model is required to predict and identify areas where the targeted study species is located or can move further if the preferable conditions and resources are available. For any living organism, climatic conditions and environmental parameters are crucial for survival in a new condition. The general ideology of the species distribution using MaxEnt is to identify the probability distribution, which has been defined during the model analysis. This distribution depends on a set of selected parameters, the occurrence of species data, and environmental characteristics suitable for the study species (Giisan et al. 2007)

This study presented the suitable habitat of *Lantana* and various biotic and abiotic factors influencing the spread of the invasion. Habitat selection by any species is based on the measure of the ecological space’s specification and resources (Basillie and Calenge 2008). Therefore in this study, the strong influence of few variables (bio18, bio8, flow accumulation, distance to road, river, elevation, soil ph) in determining the habitat selection for lantana was found to be highest. The study’s results provide the managers with valuable information on the present invasion status of *Lantana camara* in several ranges.

Past studies recorded that Lantana is mostly abundant in this region’s shrub cover in this region (Babu et al. 2009). Grazing, lopping, logging also observed during the survey in this area regularly apart from vehicle transition. Published studies express that where logging has decimated wet sclerophyll woodlands and rainforests, and gaps are made; this permits lantana to infringe on the backwoods. Further logging disturbs the infection and makes it feasible for lantana to spread or become thicker (Waterhouse and Norris 1970; Day et al. 2003). We have encountered incidents of logging, lopping and grazing within the protected area boundary during our field work. These incidents cannot be a random encounter as signs of continuous understory clearance were prominent. Thus the anthropogenic pressure in these areas can be termed as one of the critical role players in the distribution and high infestation of Lantana. Thus, the MaxEnt model prediction of high suitable areas near these zones is significant.

Previous studies in this landscape (Priyanka and Joshi 2013; Ramaswamy et al. 2016) on *Lantana* distribution highlighted areas with high invasion. The extreme invasion was reported in from Corbett TR in Jhirna, Dhikala, and Bijrani ranges. Other ranges such as Morghatti, Halduparao, Mundiyapani, Hathikund, and Kalagarh appeared to spearhead a more recent species movement in eastern and northern Kalagarh TR regions in 2013. In Rajaji National Park alone, the very high infestation was accounted for in southern and western regions like Haridwar, Dholkhand, Motichur, and Chilla range. According to the study published by Priyanka and Joshi (2013), other regions such as Chillawali, Kansrao, and Mohand have shown a more recent movement of species in northern and eastern portions of Rajaji TR.

The present study gives a current perspective of the expansion of the invaded regions in the landscape. As per the MaxEnt result prediction, Kalagarh, Bijrani, Dhikala, Jhirna, Maidavan, Palain, Kanda, Halduparao, Mandal, and Dheela holds high potential areas of *Lantana* habitat. Maidavan, Halduparao, Kalagarh ranges were reported a very recent movement of invasion previously now holds four major invasive hotspots in the region (fig 6). In Rajaji TR apart from Haridwar, Chilla, Dholkhand and Motichur, the invasion was reported previously (Priyanka and Joshi 2013; Upadhaya et al. 2018) Beribara, Gohri, Kansro and Mundal range are now invaded with *Lantana*. The middle part of the western Rajaji TR, which has high elevation showed no suitability for *Lantana* invasion. Also, in Rajaji TR areas which showed less potentiality to the invasion are mainly in high elevation. For Corbett TR, the trend is slightly different. The reason behind that is the concentration of carbon in the soil characteristics and pH level of the areas. In Rajaji TR, plain areas showed low pH value <0.5, and in Corbett the same can be seen in the high elevated areas (Maidavan, Palain, Mandal). Whereas the soil carbon content in these areas of both Rajaji and Corbett TR is high. Our model showed the 3.2% contribution of soil carbon content and in the distribution map, while soil pH has the lowest contribution, 1%. Published literature showed that heavy litter of *Lantana* canopy could escalate the amount of nitrogen and carbon in the soil, which is the crucial factor of its dense proliferation (Sharma and Raghubanshi, 2009). Expansion of the habitat suitability of *Lantan*a to the higher regions of the Himalaya was found in recent studies (Saurabh et al. 2019; Kosaka et al. 2010).

**Figure 6.**
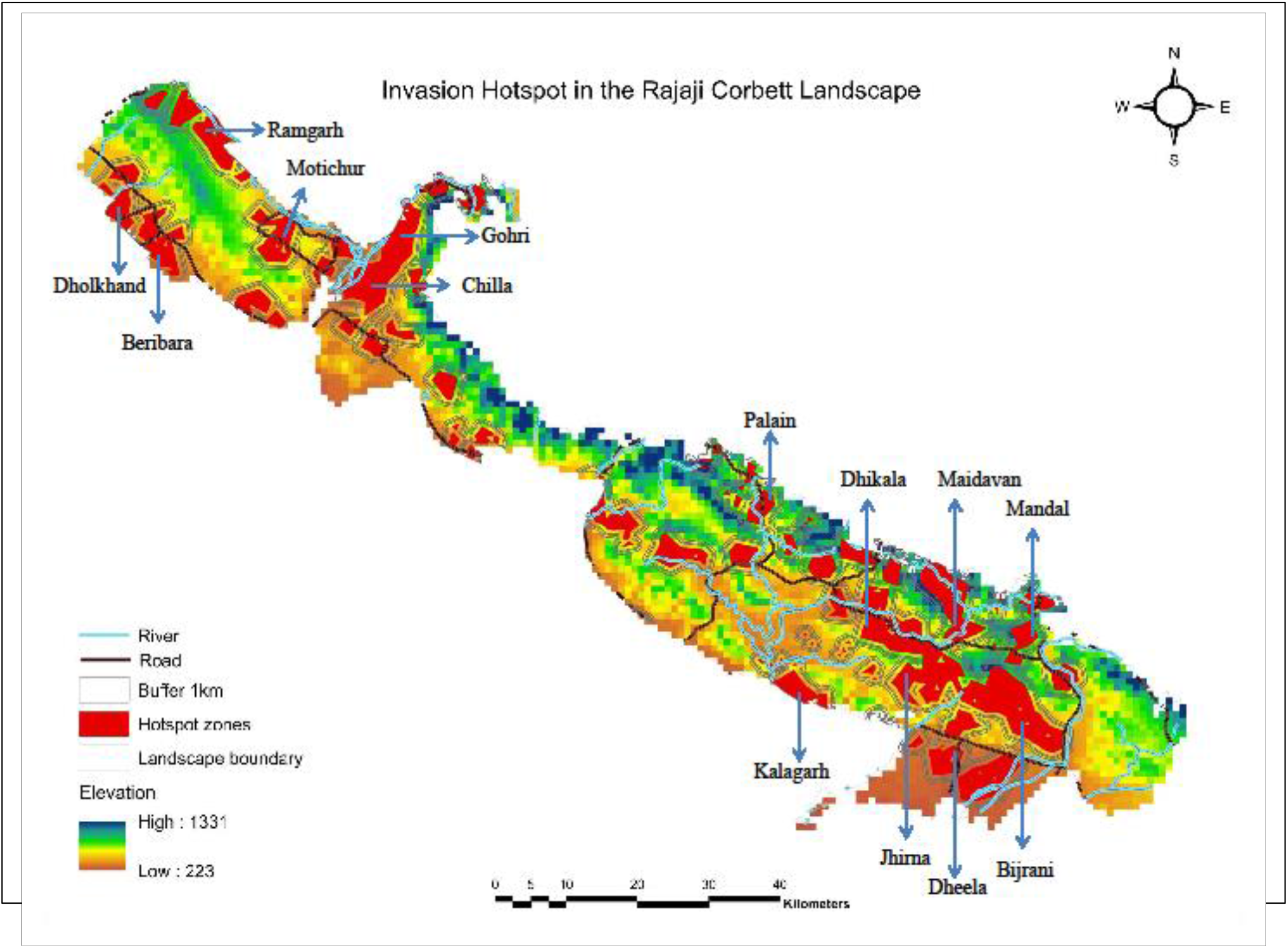
Hotspot map: showing priority sites for lantana management

Amongst the forest division, Shyampur has the high potential areas of invasion. In the lower elevation of Laldhang and Kotdwar forest division, moderate potential areas are visible. The extreme northeast part of the landscape (Kota range, fig 1) showed very low suitability of the lantana invasion. The landscape has multiple networks of streams, which are the main source of water. These streams are locally known as “Raws”. These raws are seasonal; during heavy rains, it overflows, which works as the plant seeds’ dispersal more rapidly. With the seasonal changes, the seeds get dry surfaces (river bed) to grow, propagate, and invade the open area (Ramaswami et al. 2016). These raws are often used as the demarcation boundary within the protected area ranges, beat, or compartments. However the use of these raws for regular monitoring, logistic delivery via vehicles or using as tourist tracks is a common practice during dry seasons. This day-to-day disturbance clears the ground vegetation in the nearby areas of the raw’s and enhances the chance of invasion by creating potential empty niches (Leu M et al. 2008). In our model, one of the most contributing variables is river flow accumulation (rc_accum 10.8%). The anthropogenic pressure around the raws creates suitable conditions for lantana to grow and spread rapidly (Upadhaya et al. 2018). We have recorded high infestation of lantana patches around the raws during our field documentation.

## Conclusion

The exponential growth of the invasive over the last few decades has drawn researchers and resource managers’ attention. It is incredibly crucial to know the geographical areas infested by the massive number of potentially invasive species. This study demonstrated the potential distribution of the invasive and several invasion hotspots in the landscape. *Lantana* eradication is a common practice done by the park managers and has been advised by many experts to control its spread, although the technique does not guarantee adequate efficiency and may not be able to stop long-run invasions (Swarbrick et al. 1998; Mack et al. 2000; Myers et al. 2000; Mason et al. 2006; Ramaswamy et al. 2016). Therefore strategically planned monitoring program is needed in every range housing invasion hotspot(s) and the need to prioritize regular monitoring protocols to assess invasion rates and extent remains critical. Although the eradication of *Lantana* is a logistically expensive method, and a landscape-level eradication program may seem virtually challenging, the findings from this study may aid in accelerating monitoring initiatives based on hotspots and patterns of invasion. First-hand invasive science information provided by the current study, may serve as crucial foundations to assist and develop a cost-efficient management strategy.

**Table 1 :** List of variables used for the analysis

| Predictor variables | Description | Scale used | Source |
| --- | --- | --- | --- |
| BIO2 | Mean Diurnal Range (Mean of monthly (max temp - min temp)) | 1km | www.worldclim.org |
| BIO3 | Isothermality (BIO2/BIO7) ($\times 100$) | 1km | www.worldclim.org |
| BIO6 | Min Temperature of Coldest Month | 1km | www.worldclim.org |
| BIO8 | Mean Temperature of Wettest Quarter | 1km | www.worldclim.org |
| BIO15 | Precipitation Seasonality (Coefficient of Variation) | 1km | www.worldclim.org |
| BIO17 | Precipitation of Driest Quarter | 1km | www.worldclim.org |
| BIO18 | Precipitation of Warmest Quarter | 1km | www.worldclim.org |
| BIO19 | Precipitation of Coldest Quarter | 1km | www.worldclim.org |
| BIO13 | Precipitation of Wettest Month | 1km | www.worldclim.org |
| Slope | Topographic characteristic | 1km | Digital elevation map.diva gis |
| Aspect | Topographic characteristic | 1km | Digital elevation map.diva gis |
| Elevation | Topographic characteristic | 1km | Digital elevation map.diva gis |
| LULC | Current land use land cover of the landscape | 1km | Modislandcover.usgs.org |
| Flow accumulation | Streams and major river flow in the landscape | 1km | Calculated using digital elevation map |
| Soil ph content at 0cm | A measure of the **acidity** or alkalinity of the **soil** | 1km | Openlandmap.org |
| Soil carbon content at 0cm | Solid terrestrial matter stored in **soils** | 1km | Openlandmap.org |

**Table 2:**
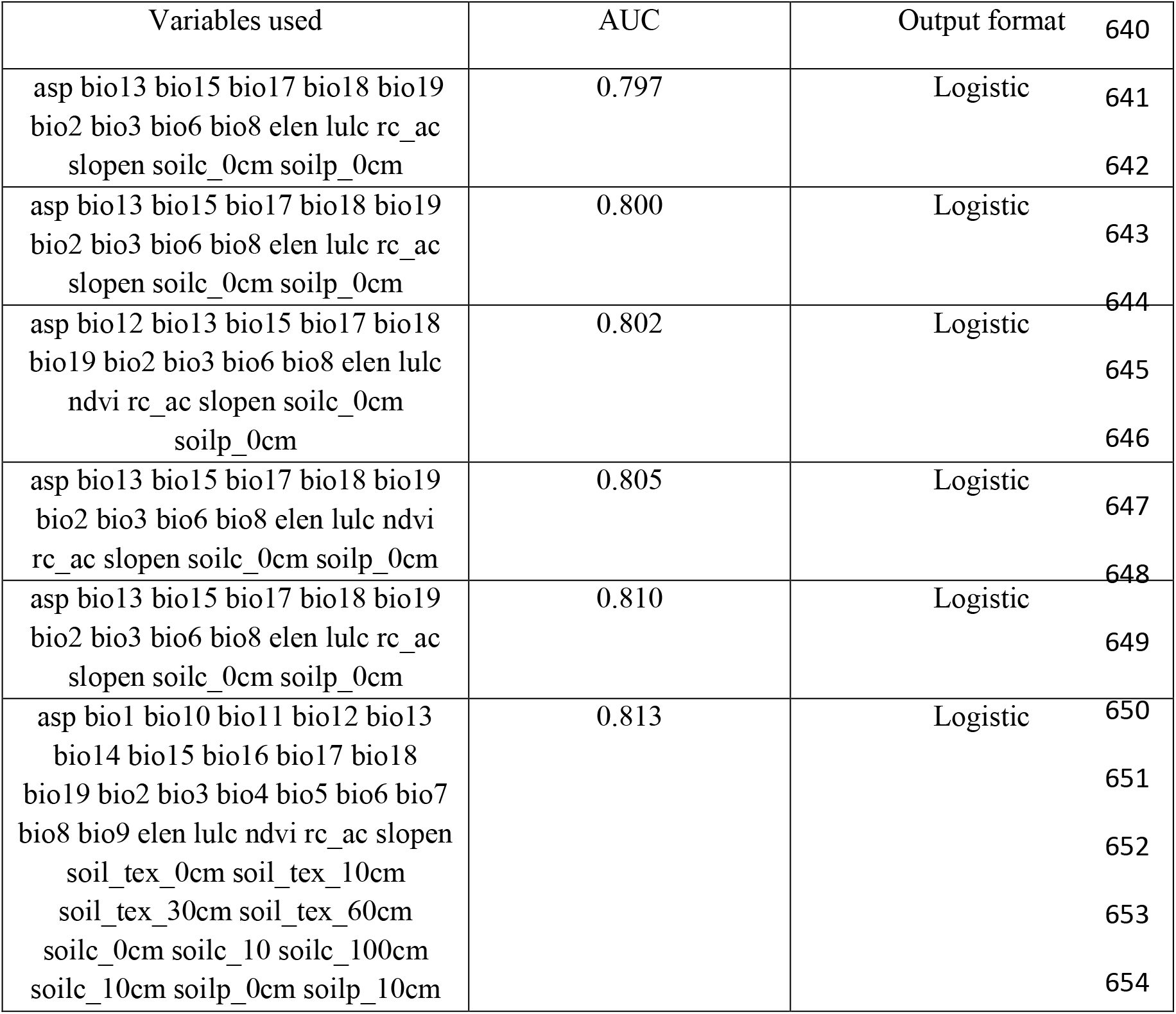
List of different models with AICC value

| Variables used | AUC | Output format | 640 |
| --- | --- | --- | --- |
| asp bio13 bio15 bio17 bio18 bio19<br>bio2 bio3 bio6 bio8 elen lulc rc_ac<br>slopen soilc_0cm soilp_0cm | 0.797 | Logistic | 641<br>642 |
| asp bio13 bio15 bio17 bio18 bio19<br>bio2 bio3 bio6 bio8 elen lulc rc_ac<br>slopen soilc_0cm soilp_0cm | 0.800 | Logistic | 643<br>644 |
| asp bio12 bio13 bio15 bio17 bio18<br>bio19 bio2 bio3 bio6 bio8 elen lulc<br>ndvi rc_ac slopen soilc_0cm<br>soilp_0cm | 0.802 | Logistic | 645<br>646 |
| asp bio13 bio15 bio17 bio18 bio19<br>bio2 bio3 bio6 bio8 elen lulc ndvi<br>rc_ac slopen soilc_0cm soilp_0cm | 0.805 | Logistic | 647<br>648 |
| asp bio13 bio15 bio17 bio18 bio19<br>bio2 bio3 bio6 bio8 elen lulc rc_ac<br>slopen soilc_0cm soilp_0cm | 0.810 | Logistic | 649 |
| asp bio1 bio10 bio11 bio12 bio13<br>bio14 bio15 bio16 bio17 bio18<br>bio19 bio2 bio3 bio4 bio5 bio6 bio7<br>bio8 bio9 elen lulc ndvi rc_ac slopen<br>soil_tex_0cm soil_tex_10cm<br>soil_tex_30cm soil_tex_60cm<br>soilc_0cm soilc_10 soilc_100cm<br>soilc_10cm soilp_0cm soilp_10cm | 0.813 | Logistic | 650<br>651<br>652<br>653<br>654 |

## Acknowledgement

The authors are grateful to the National Tiger Conservation Authority for funding this study under the grant (No. WII/KR/PROJECT/DRONE/2015/018) and Dr.V.B. Mathur (former Director) and Dr. G.S. Rawat (former Dean) of Wildlife Institute of India for facilitating the same. Soumya Dasgupta is thanked for his inputs towards designing the field survey and crystalizing the statistical framework. Chief Wildlife Warden Uttarakhand, Field Director Rajaji Tiger reserve and Field Director Corbett Tiger Reserve are thanked for providing permits to undertake surveys in protected areas. Mr Shehzad and Sipu are thanked for assistance in field.

